# *Gardnerella fastidiominuta* sp. nov. isolated from the female urinary microbiome

**DOI:** 10.64898/2026.03.30.715431

**Authors:** Liliane Pinto Ferrador, Bárbara Duarte, Filipa Grosso, Teresa Gonçalves Ribeiro, Luísa Peixe

**Affiliations:** UCIBIO, i4HB Faculdade de Farmácia, Universidade do Porto, Rua de Jorge Viterbo Ferreira 228, 4050-313 Porto, Portugal; Culture Collection of Porto (CCP), Faculdade de Farmácia, Universidade do Porto, Rua de Jorge Viterbo Ferreira 228, 4050-313 Porto, Portugal

**Keywords:** *Gardnerella*, novel species, urinary microbiome, polyphasic taxonomy, phylogenomics, genome-based taxonomy

## Abstract

The genus *Gardnerella* comprises a group of fastidious bacteria associated with the female urogenital tract and has undergone extensive taxonomic revision in recent years. In this study, a bacterial strain, designated CCPDSMᵀ, was isolated from the female urinary microbiome and subjected to a comprehensive polyphasic taxonomic characterization. The 16S rRNA gene sequence confirmed that this strain is a member of the genus *Gardnerella*, and phylogenetic analyses based on *cpn60* sequences, together with phylogenomic reconstruction placed strain CCPDSMᵀ within the genus *Gardnerella* as a distinct and well-supported lineage.

Genome-based relatedness indices (ANIb, ANIm, TETRA and dDDH), demonstrated clear separation of CCPDSMᵀ from all validly published *Gardnerella* species. In contrast, comparisons with two publicly available closely related genomes yielded values above accepted species delineation thresholds, supporting their assignment to the same taxon. Phenotypic characterization, together with genome-based functional predictions, revealed a fastidious, fermentative metabolic profile that further differentiated CCPDSMᵀ from its closest relatives, while remaining consistent with traits characteristic of the genus.

On the basis of combined phylogenetic, genomic and phenotypic evidence, strain CCPDSMᵀ is proposed as representing a novel species within the genus *Gardnerella*, for which the name *Gardnerella fastidiominuta* sp. nov. is proposed, with strain CCPDSMᵀ (=CECT 31324ᵀ=CCP 588ᵀ) designated as the type strain. This study expands the recognized diversity of *Gardnerella* and highlights the female urinary tract as a reservoir of previously uncharacterized species within this genus.

## Introduction

The genus *Gardnerella* belongs to the phylum *Actinobacteria*, class *Actinomycetia*, order *Bifidobacteriales*, and family *Bifidobacteriaceae* (1). For nearly seven decades, *Gardnerella vaginalis* was regarded as the sole representative of the genus and was primarily associated with bacterial vaginosis, one of the most prevalent vaginal dysbioses in women (2).

This long-standing taxonomic simplicity was substantially revised in 2019, following the description of three additional species (*Gardnerella leopoldii*, *Gardnerella piotii*, and *Gardnerella swidsinskii*), supported by integrative genomic, proteomic, and phenotypic analyses, together with the recognition of several genomic species (3). Since the initial taxonomic expansion of the genus, additional *Gardnerella* species have been described (i.e., *Gardnerella pickettii, Gardnerella greenwoodii, Gardnerella massiliensis*, *Gardnerella bretellae*, *Gardnerella phocaensis*), further underscoring the still incompletely resolved diversity within this genus (4, 5).

Members of the genus *Gardnerella* are Gram-variable, facultatively anaerobic bacteria commonly detected in the female urogenital tract, although their ecological roles and clinical relevance remain incompletely understood (6). Historical taxonomic constraints, whereby isolates were largely assigned to *G. vaginalis* or poorly resolved genomic groups, have hindered accurate species-level interpretation of their distribution, ecological niches and potential functional differentiation.

Increasing evidence indicates that multiple *Gardnerella* species are detected in the urogenital tract of asymptomatic women (7, 8), often at low relative abundance and frequently in microbiota communities depleted of *Lactobacillus* spp. (9, 10). These observations underscore the need for continued refinement of *Gardnerella* taxonomy to support robust ecological and clinical studies (11). Although *Gardnerella* species have been primarily characterized from vaginal isolates (3), their presence in the female urinary tract has become increasingly recognised using culture-dependent and independent approaches (12, 13). Nevertheless, the diversity of *Gardnerella* species inhabiting this niche remains incompletely characterised.

During an ongoing investigation of the female urinary microbiome in our laboratory (9, 12), a bacterial strain representing a putative novel species within the *Gardnerella* genus was isolated from a urine sample obtained from an asymptomatic postmenopausal woman. Comparative phylogenetic, genomic and phenotypic analysis demonstrated that this strain represents a distinct and previously unrecognised species within the genus. Based on a polyphasic approach, we propose the name *Gardnerella fastidiominuta* sp. nov. for this novel species, with strain CCPDSM^T^ designated as the type strain.

## Methods

### Sample processing and bacterial strain isolation

A urine sample obtained from an asymptomatic postmenopausal woman was processed as previously described (11). Briefly, 100 µL of urine was plated onto Columbia agar supplemented with 5% sheep blood (Biogerm, Portugal), and HiCrome UTI agar (HiMedia, India). Plates were incubated at 37 °C for 48 h, under aerobic, microaerophilic, and anaerobic conditions. Colonies displaying distinct morphotypes were selected, subcultured to purity, and stored at -80 °C in tryptic soy broth (Liofilchem, Italy) supplemented with 20% (v/v) glycerol.

### Morphological, physiological and biochemical tests

Growth was assessed at 20, 25, 30, 37, 40 and 45 °C on Columbia agar supplemented with 5 % sheep blood (Frilabo, Portugal) and chocolate agar (Liofilchem, Italy) after 24, 48, and 72 h of incubation, under different atmospheric conditions (aerobic, microaerophilic, and anaerobic). Growth at different pH values (4.5, 5.5, 7.5, 8.0, and 9.0) and NaCl concentrations [0.5 %, 1.0 %, and 1.5 %, (w/v)] was evaluated in Brain Heart Infusion broth supplemented (BHIs) with starch (0.1%), yeast extract (0.5%), gelatin (2%), and glucose (0.1%), after 24, 48, and 72 h of incubation at 37°C, under anaerobic conditions using a Synergy HT 96-well microplate reader (Biotek, USA).

Colony morphology was examined under optimal growth conditions. Gram staining was carried out using a commercial Gram-staining kit (bioMérieux, France). Biochemical characterisation was performed using API® CORYNE and API® ZYM systems (bioMérieux, France), following the manufacturer’s instructions. β-galactosidase activity was tested using ONPG tablets (Sigma Aldrich, USA), and oxidase activity was determined using oxidase reagent strips (ThermoFisher, USA), according to the manufacturer’s recommendations.

### Genome sequencing and analyses

Genomic DNA was extracted from culture grown in 10 ml BHIs for 48 h at 37 °C, under anaerobic conditions. Cells were harvested by centrifugation at 13,000 rpm for 10 min, and genomic DNA was extracted using the GenFind V3 DNA Extraction Kit (Beckman Coulter, USA). DNA concentration was assessed by fluorometry using a Qubit™ 4 fluorometer (Thermo Fisher Scientific).

Whole-genome sequencing was performed on an Illumina NovaSeq 6000 platform (2×150 bp paired-end reads). Initial bioinformatic analyses were carried out using the Bacterial and Viral Bioinformatics Resource Center (BV-BRC) platform (14). Raw reads were trimmed with Trim Galore 0.6.5 (15) (http://www.bioinformatics.babraham.ac.uk/projects/trim_galore/), and quality was checked with FastQC 0.11.9 (www.bioinformatics.babraham.ac.uk/projects/fastqc/). *De novo* genome assembly was performed using Unicycler v0.4.8 (16), and assembly quality was assessed with QUAST v5.0.2 (17). Genome completeness and contamination were assessed using CheckM (18). Genome annotation was performed using the NCBI Prokaryotic Genome Annotation Pipeline and the RAST toolkit (RASTtk) (19, 20).

### Data collection

Publicly available *Gardnerella* genomes were retrieved from the NCBI database to ensure representative coverage of the genomic diversity within the genus. This dataset included genomes of type strains of all validly published *Gardnerella* species, as well as non-type strain genomes (up to 4 genomes/species) previously assigned to *Gardnerella* species and genomic species based on curated classifications proposed by Vaneechoutte *et al*. (2019) and Sousa *et al*. (2023) (3,4). From the selected genomes, complete or near-complete sequences of the 16S rRNA gene and *cpn60* were extracted for downstream phylogenetic analyses.

In addition, the Similar Genome Finder tool implemented in the BV-BRC platform (https://www.bv-brc.org/app/GenomeDistance) was used to identify genomes showing high genomic similarity to strain CCPDSM^T^, in order to assess whether closely related genomes potentially belonging to the same species were present in public databases. Genomes identified by this approach were subsequently retrieved for comparative genomic analyses.

### Genomic relatedness analyses

Average nucleotide identity based on BLAST+ (ANIb) or MUMmer (ANIm), as well as tetranucleotide signature correlation index (TETRA) scores, were calculated using JSpeciesWS (21). In addition, digital DNA-DNA hybridization (dDDH) values were determined using the TYGS web server, following the recommended Formula 2 (22, 23). Phylogenomic analysis was conducted using the BV-BRC Bacterial Genome Tree Service (Codon Tree method) and the default parameters. The method selects single-copy BV-BRC global protein families (PGFams), builds concatenated alignments, and infers a maximum-likelihood tree using RAxML with bootstrap support (24). The resulting tree was visualised and annotated using iTOL v6 (25).

Clustered Regularly Interspaced Short Palindromic Repeats and CRISPR-associated (Cas) genes were identified using CRISPRcasFinder (26). Prophage regions were identified using PHASTER (27). Antimicrobial resistance genes and plasmid replicons were investigated using CARD, ResFinder 4.7.2, and PlasmidFinder 2.0.1, respectively (28–30). BAGEL4 was used to mine bacteriocin gene clusters (31).

### Phylogenetic analysis

Complete nucleotide sequences of the 16S rRNA gene and *cpn60* were extracted from the genome of strain CCPDSM^T^, type strains of *Gardnerella* species, and closely related genomic species, using UniPro UGENE v.50 (32) or retrieved from the GenBank database (33). Sequence alignments and similarity analysis were generated using MEGA version 7.0 (http://www.megasoftware.net/) (34). Phylogenetic trees were constructed using the neighbour-joining method (35), and genetic distances were estimated under the Kimura two-parameter model (36). The reliability of internal branches was assessed from bootstrapping based on 1000 resamplings (37).

## Results and Discussion

### Phylogenetic analysis based on 16S rRNA gene and *cpn60*

Analysis of the 16S rRNA gene sequence confirmed that strain CCPDSM^T^ belongs to the genus *Gardnerella*, with the closest relatives being *Gardnerella* genomic species 10 strain 1500E and *G. swidsinskii* strains GS 10234 and JNFY3, showing sequence identities of 99.93% and 99.89%, respectively (Fig. 1). This high level of sequence identity is consistent with previous studies demonstrating that closely related *Gardnerella* species often share nearly identical 16S rRNA gene sequences (4), thereby limiting the discriminatory power of this marker within the genus.

**Fig. 1:**
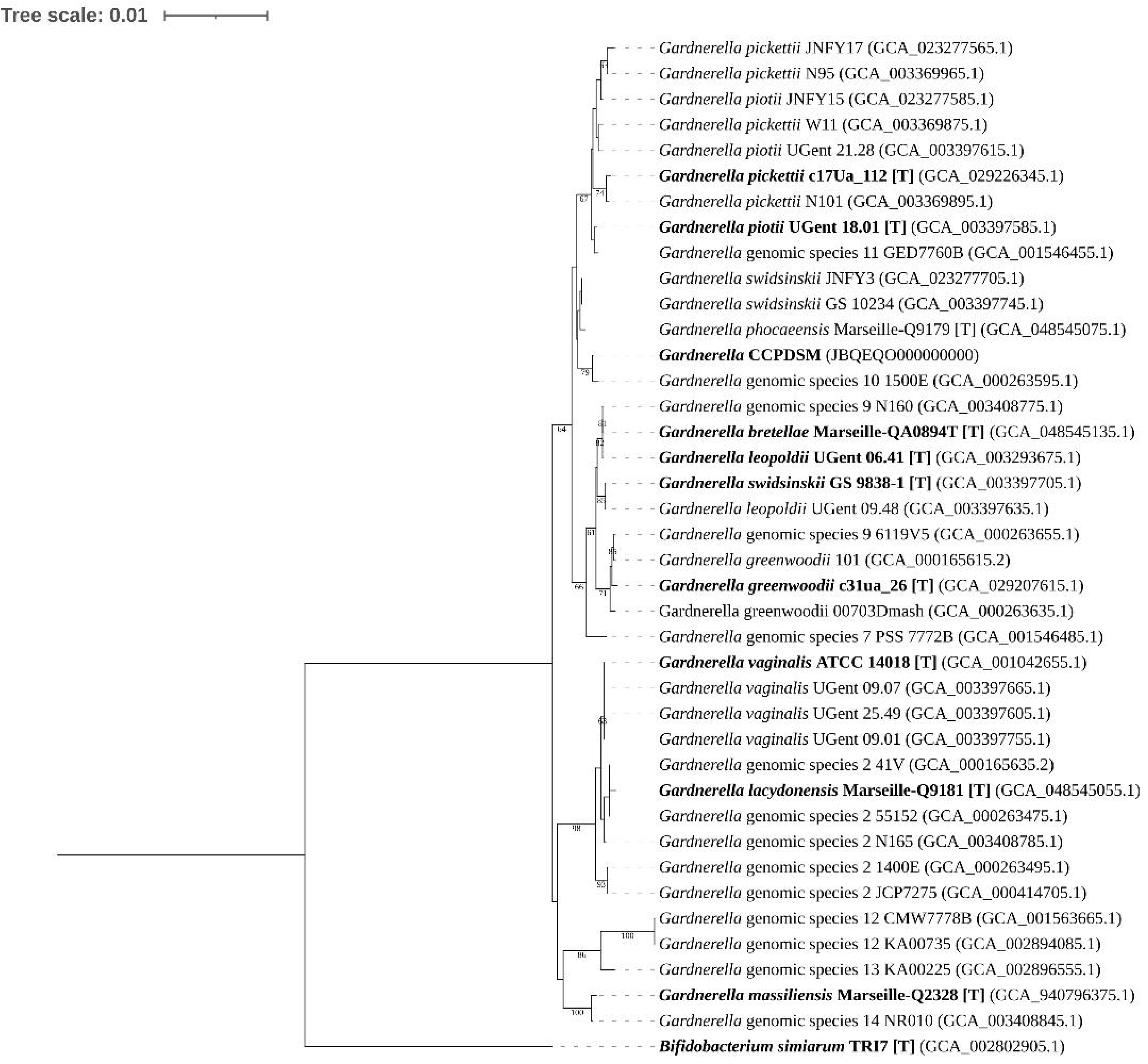
Neighbour-joining tree based on complete 16S rRNA gene sequences (1538 bp) showing the phylogenetic relationships of strain CCPDSM^T^ (boldface type), type strains of the genus *Gardnerella* (boldface type denoted by “[T]” at the end of the strain name), and closely related genomic species. *Bifidobacterium simiarum* was used as the outgroup. Bootstrap percentages (based on 1000 replications) are shown at nodes. Only values above 60% are shown. Bar, 0.02 substitutions per nucleotide position. The available accession numbers of the sequences used are indicated in parentheses. Strains JCP8481B and JCP8481A, belonging to Gardnerella genomic species 7, were not included due to incomplete 16S rRNA gene sequences.

In contrast, phylogenetic analysis based on the *cpn60* provided higher resolution and demonstrated that strain CCPDSM^T^ did not cluster with any currently validly published *Gardnerella* species or previously defined genomic species (Fig. 2). The highest identity was observed with *Gardnerella greenwoodii* strain 00703Dmash (90.2%), followed by *G. greenwoodii* strain 101 (90.0%). These values are well below accepted species-thresholds for *cpn60* (4), supporting that strain CCPDSM^T^ constitutes a distinct phylogenetic lineage within the genus *Gardnerella*.

**Fig. 2:**
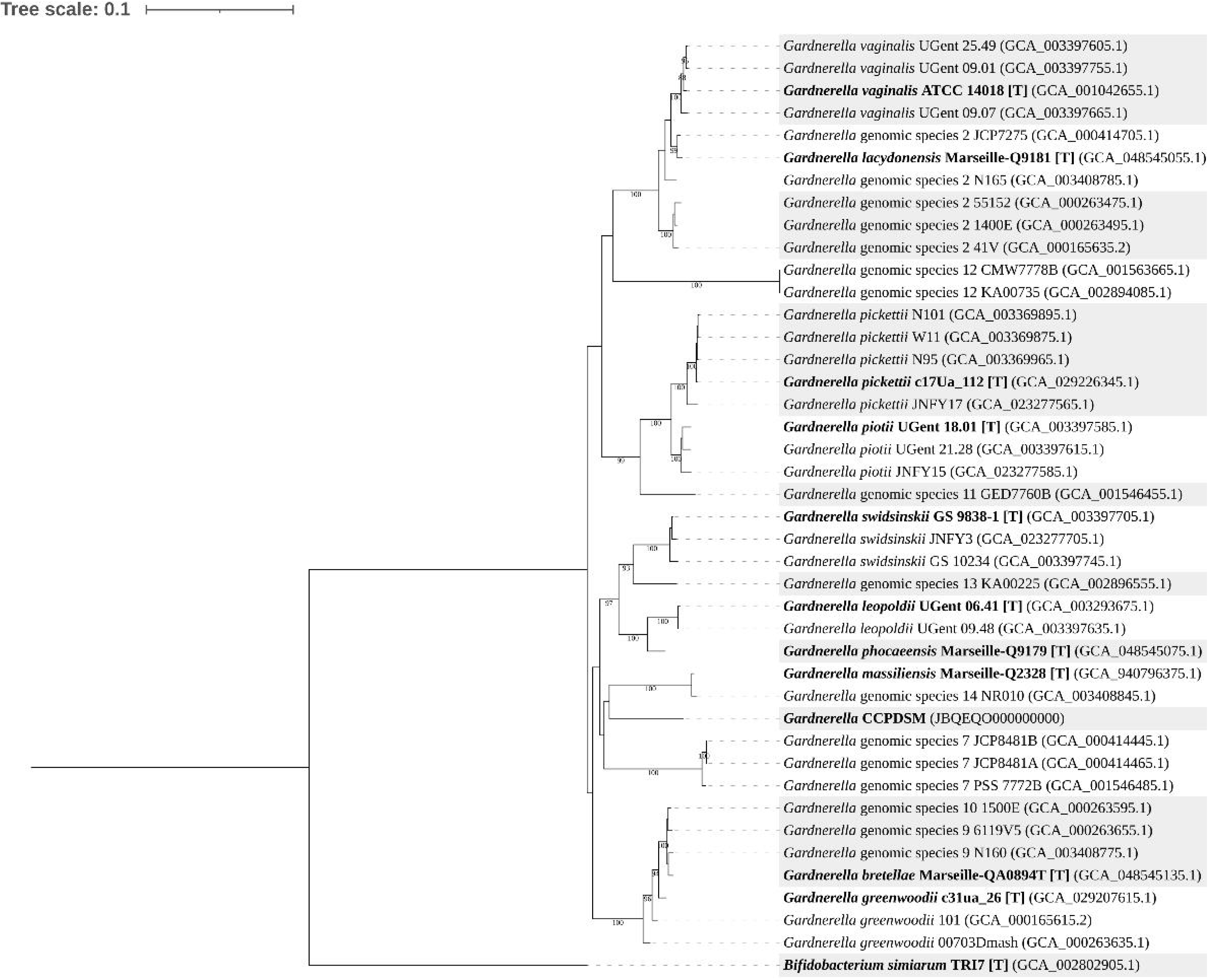
Neighbour-joining tree based on complete *cpn60*-gene sequences (1626 bp) showing the phylogenetic relationships of strain CCPDSM^T^ (boldface type), type strains of the genus *Gardnerella* (boldface type denoted by “[T]” at the end of the strain name), and closely related genomic species. *Bifidobacterium simiarum* was used as the outgroup. Bootstrap percentages (based on 1000 replications) are shown at nodes. Only values above 80% are shown. Bar, 0.02 substitutions per nucleotide position. The available accession numbers of the sequences used are indicated in parentheses.

### Genome-based features, relatedness and phylogenomic inference

The draft genome of strain CCPDSM^T^ comprised 1,645,963 bp distributed across 96 contigs. Genome quality assessment indicated a completeness of 99.8% and a contamination rate of 0.2%, supporting the suitability of the assembly for comparative genomic analyses. The genomic G+C content was 43.35%, and the N50 value was 203,116 bp. Genome annotation identified 1,333 genes in total, including 1,282 CDSs (1,275 protein-coding CDSs with functional assignment), 3 rRNA genes and 45 tRNA genes. These genomic features fall within the range reported for members of the genus *Gardnerella* (Table 1), supporting the assignment of strain CCPDSM^T^ to this genus.

**Table 1.**
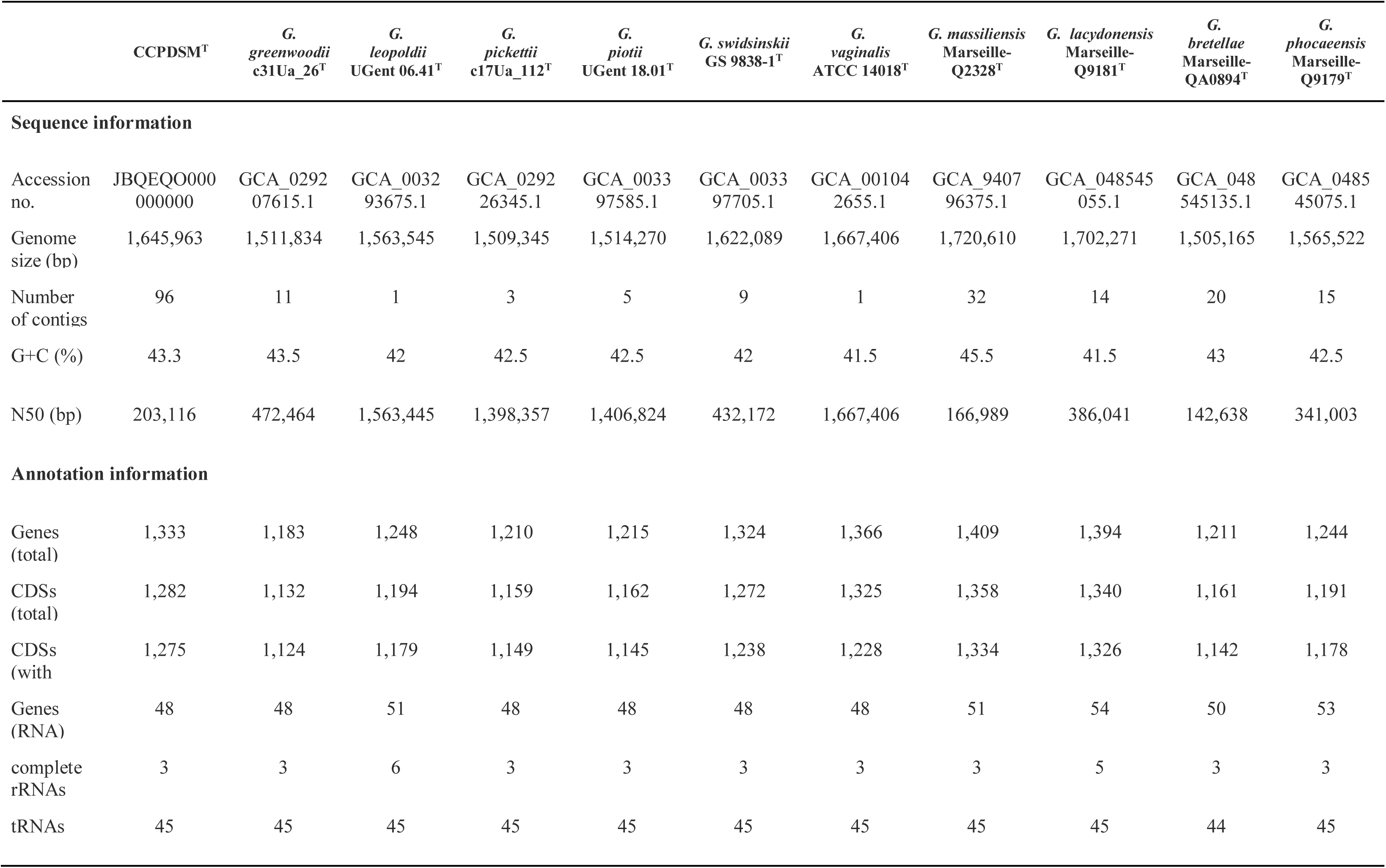
Genomic features of strain CCPDSM^T^ and type strains of *Gardnerella* species.

Similarity searches performed with the BV-BRC Similar Genome Finder identified two publicly available genomes with high genomic similarity to strain CCPDSMᵀ, each sharing more than 315 of 1000 MinHash k-mers and exhibiting low Mash distances (≤ 0.035). These genomes correspond to metagenome-assembled genomes (MAGs) recovered from human vaginal samples (SRR16916876_bin.13_metaWRAP_v1.3_MAG, assembly accession GCA_946997395.1, USA, 2004, completeness 56.2%, contamination 4.8%; and SRR17635500_bin.3_metaWRAP_v1.3_MAG, assembly accession GCA_947253005.1, USA, 2008, completeness 60.5%, contamination 1.1%). Genomic relatedness indices between CCPDSMᵀ and these MAGs further supported a close relationship (Table 2): ANIb values were ≥ 95.77%; ANIm values were ≥ 96.26%; TETRA values exceeded 0.990; and dDDH values were 65.6% for SRR16916876_bin.13_metaWRAP_v1.3_MAG and 73.1% for SRR17635500_bin.3_metaWRAP_v1.3_MAG, with the former falling slightly below the conventional 70% species delineation threshold (38).

**Table 2.**
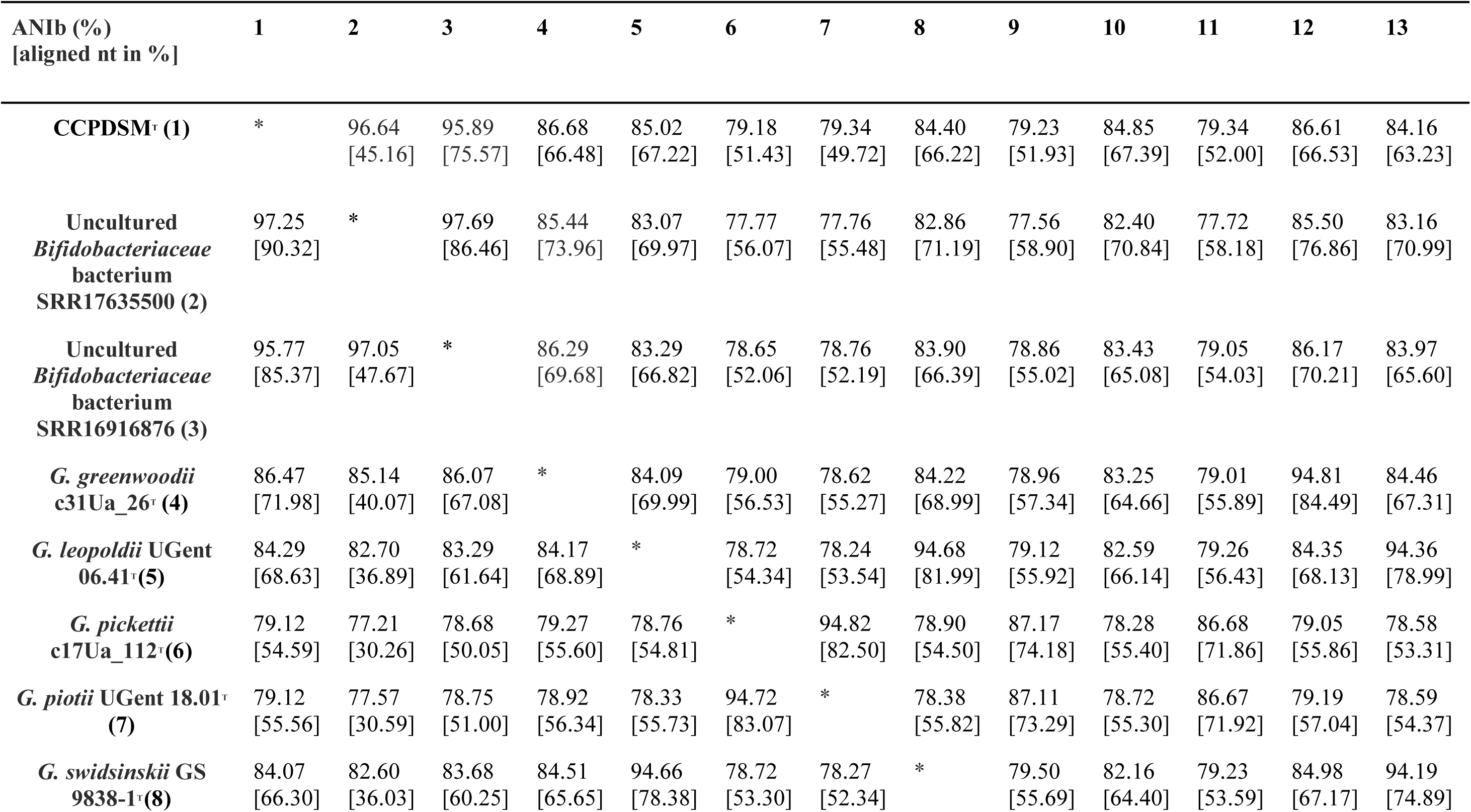
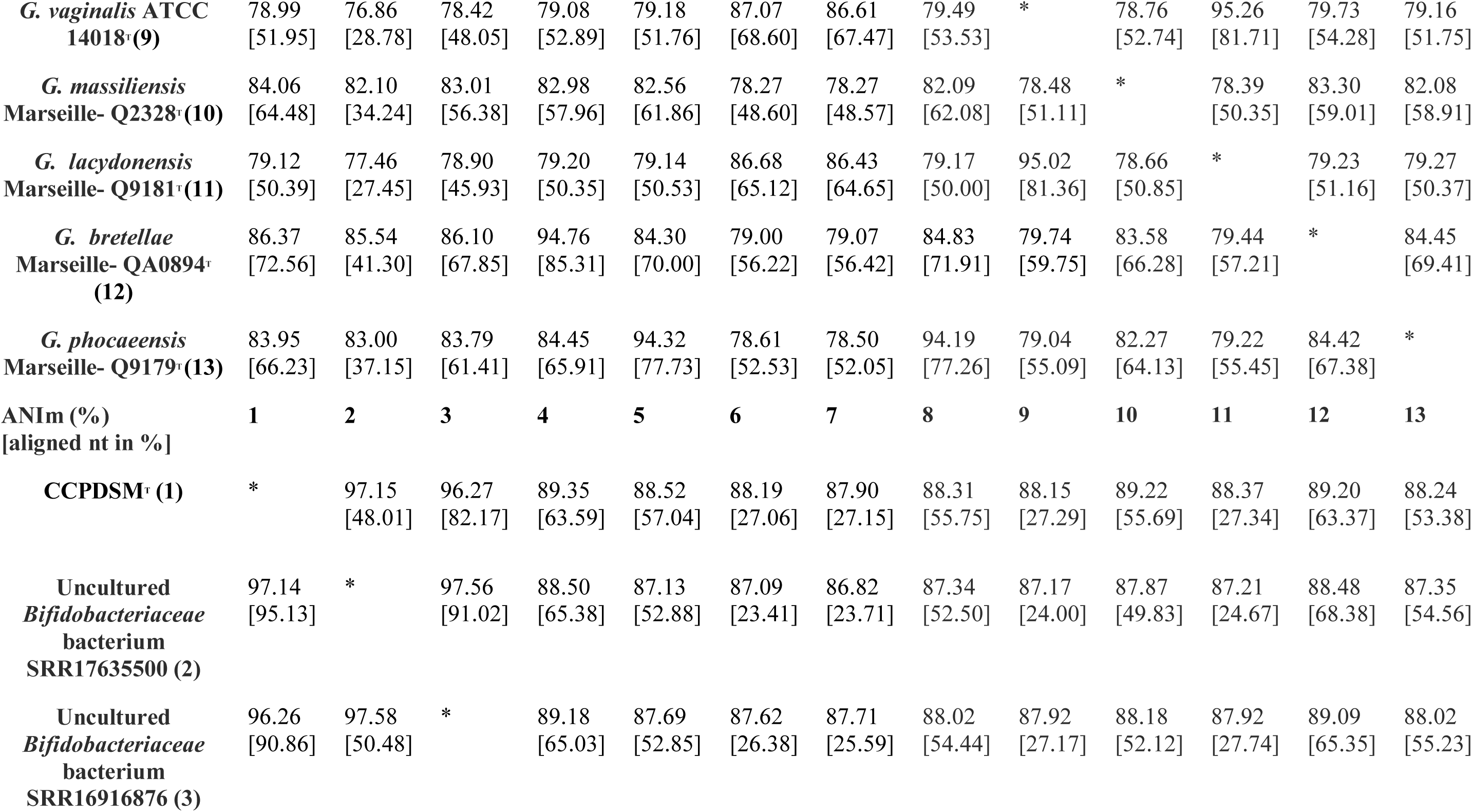
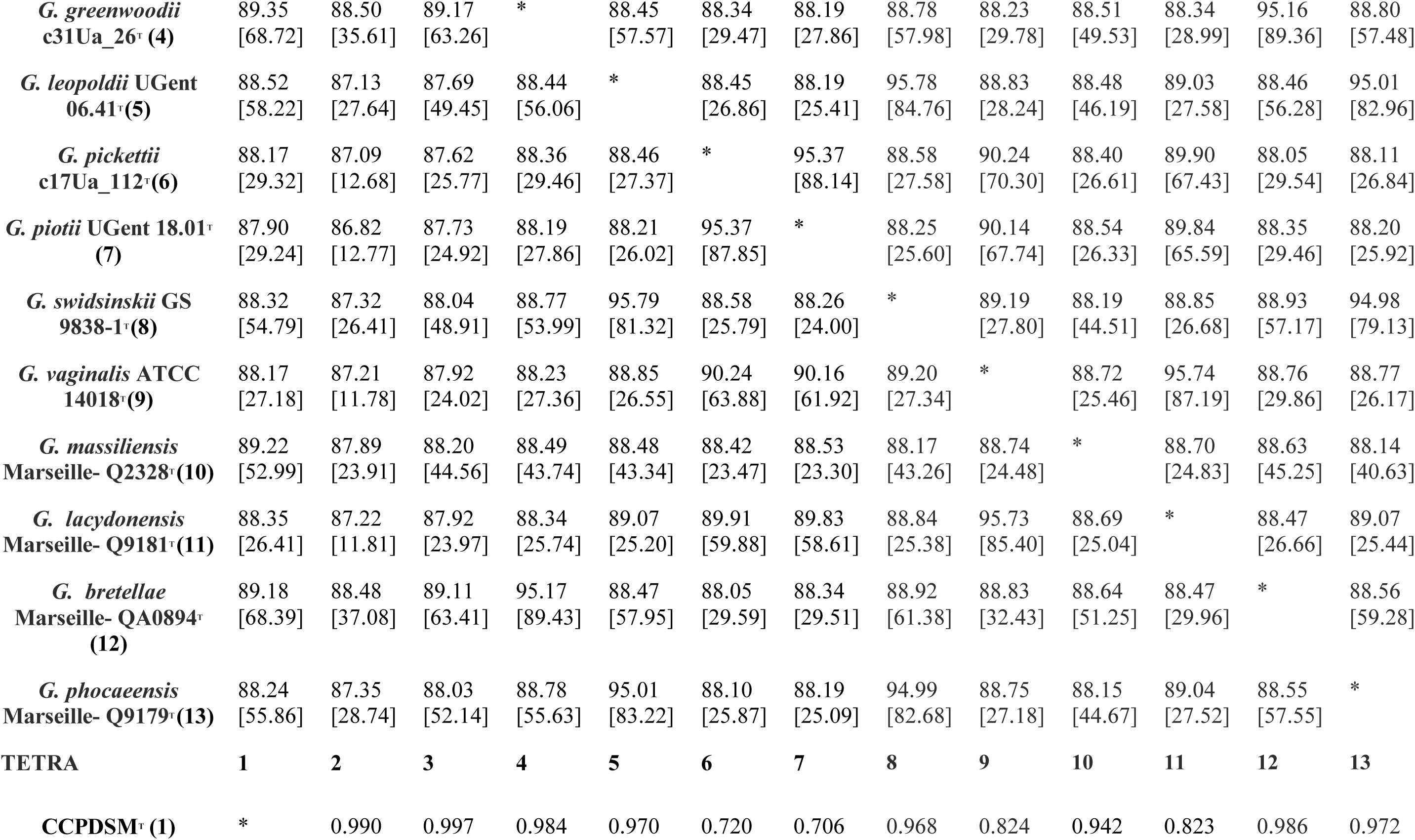
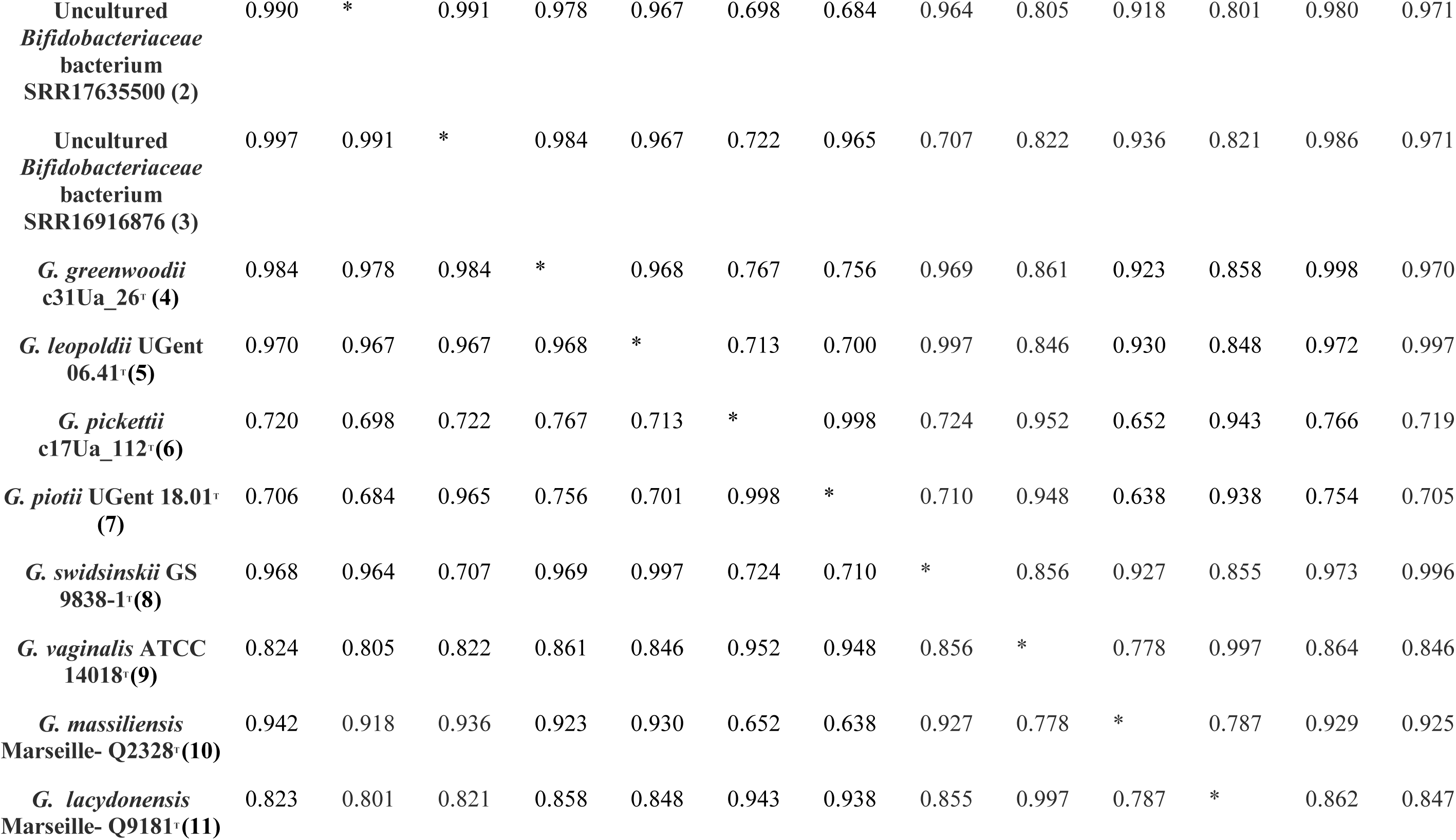
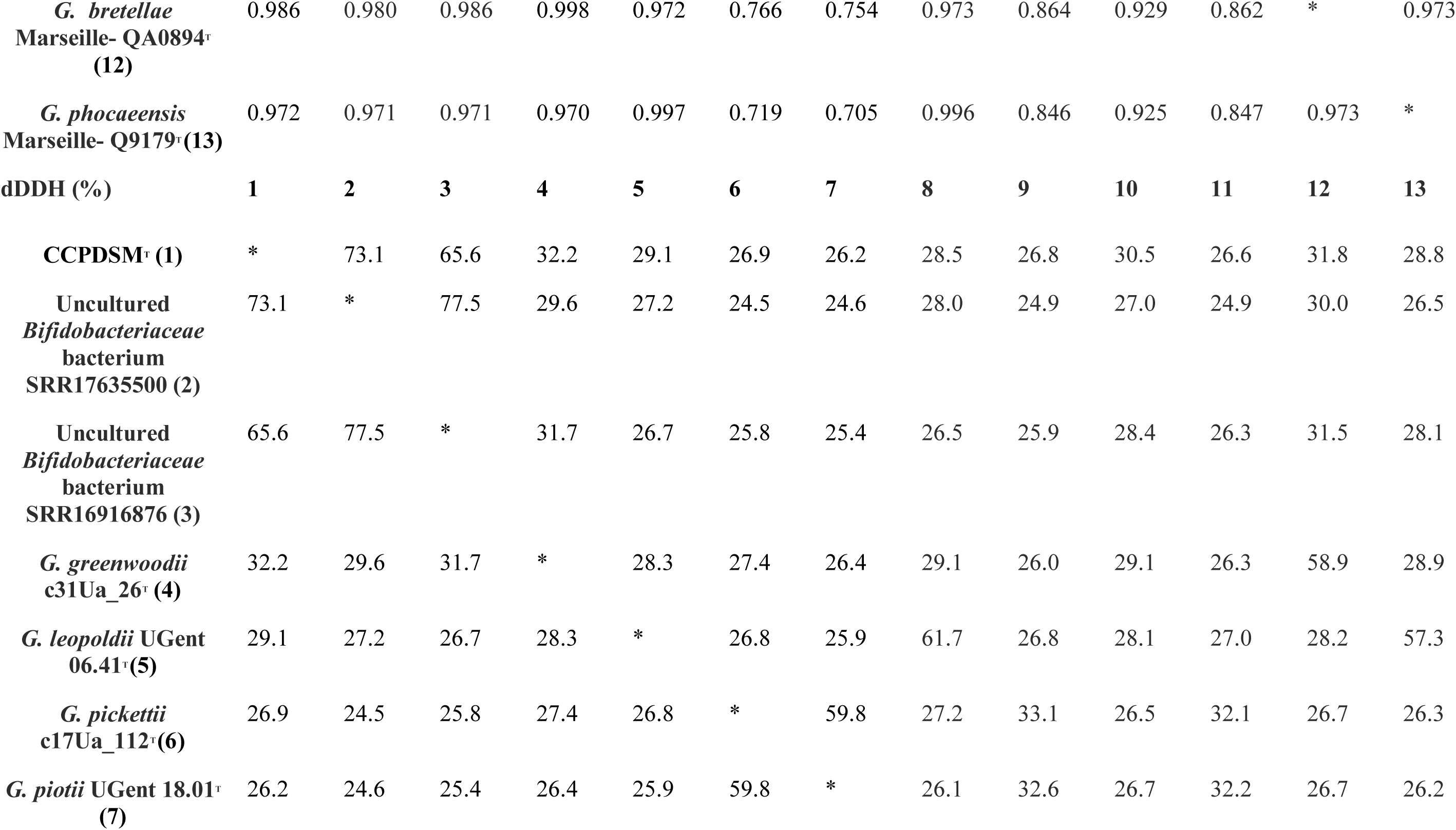
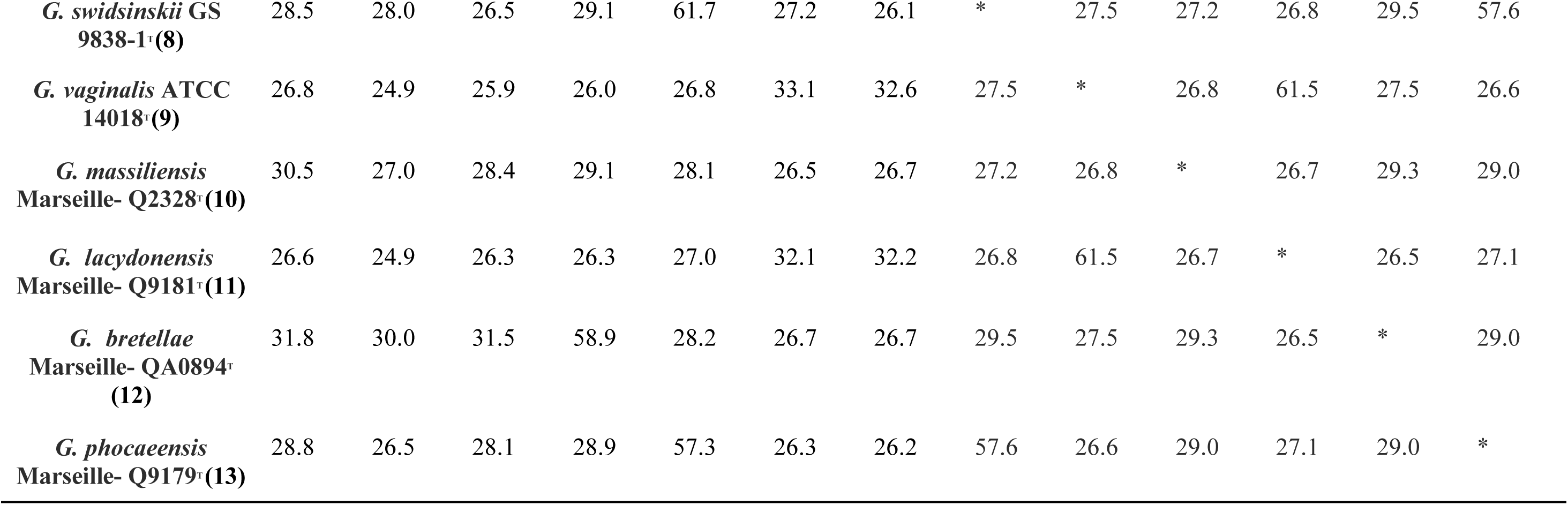
Pairwise genomic relatedness indices (ANIb, ANIm, TETRA and dDDH) between strain CCPDSM^T^ and type strains of *Gardnerell*a species.

Taken together, the ANIb/ANIm, TETRA, Mash, and MinHash metrics suggest that both MAGs likely belong to the same species-level lineage as strain CCPDSMᵀ (40, 41), although their incomplete nature precludes definitive assignment

To evaluate genomic relatedness at the species level across the genus, genome relatedness indices were calculated between strain CCPDSMᵀ and the type strains of all validly published *Gardnerella* species (Table 2). ANIb values ranged from 79.18% to 86.68%, all below the generally accepted species threshold (40). The highest ANIb value was observed between CCPDSM^T^ and *G. greenwoodii* c31Ua_26^T^ (86.68%), while ANIm values showed a similar pattern, with a maximum of 89.35% between CCPDSM^T^ and *G. greenwoodii* c31Ua_26^T^ (41). TETRA values were below 0.986, lower than values typically observed among strains of the same species (40). In addition, dDDH values ranged from 26.2% to 32.2%, with the highest value observed against *G. greenwoodii* c31Ua_26^T^, and all well below the 70% species delineation threshold (38).

Phylogenomic reconstruction based on concatenated alignments of 100 conserved single-copy core protein families positioned strain CCPDSM^T^ within the genus *Gardnerella* as a distinct, and well-supported lineage (Fig. 3). Strain CCPDSM^T^, together with the two metagenome-assembled genomes (SRR16916876_bin.13_metaWRAP_v1.3_MAG and SRR17635500_bin.3_metaWRAP_v1.3_MAG), formed a monophyletic clade with strong bootstrap support (100%), clearly separated from all other *Gardnerella* species or genomic species.

**Fig. 3:**
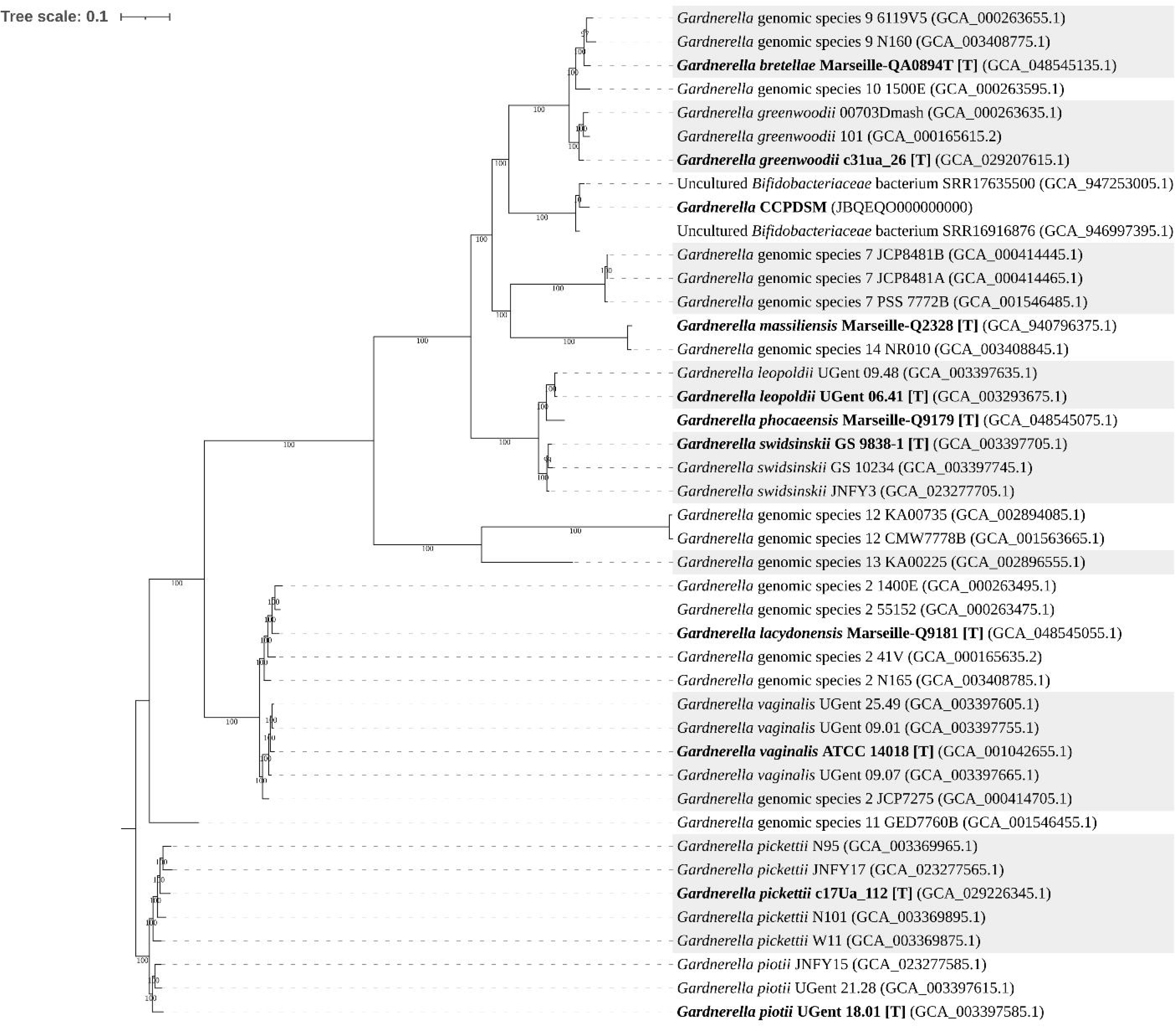
Phylogenomic tree of the genus *Gardnerella*. Phylogenomic tree based on concatenated alignments of conserved single-copy protein families, reconstructed using the BV-BRC Bacterial Genome Tree service. The tree shows the position of strain CCPDSM^T^ within the genus *Gardnerella*. Bootstrap support values above 70% are indicated at branch nodes. Type strains are boldfaced and marked with a “[T]” at the end of the strain name. The available accession numbers of the sequences used are indicated in parentheses.

The overall phylogenomic topology was congruent with genome-based relatedness indices (ANIb, ANIm, TETRA, and dDDH), and with *cpn60*-based phylogenetic analyses, providing consistent and robust evidence for species-level separation of strain CCPDSM^T^ within the genus *Gardnerella*.

### Phenotypic characteristics

Cells of strain CCPDSM^T^ are Gram-staining-positive, non-spore-forming, non-motile coccobacilli. Growth occurs under anaerobic and microaerophilic conditions on Columbia agar supplemented with 5% sheep blood and chocolate agar, after 48-72h of incubation, at temperatures between 25 and 40°C (optimum 37°C). No growth is observed under aerobic conditions. The strain grows within a pH range of 5.5–7.5 and at NaCl concentrations of 0.5–1.0%. Colonies are pinpoint, very small, circular, opaque, white to greyish, and non-haemolytic on blood agar.

Strain CCPDSM^T^ is catalase- and oxidase-negative, and does not exhibit β-galactosidase activity (ONPG test negative). API ® Coryne and API® ZYM assays revealed positive reactions for maltose, leucine arylamidase, acid phosphatase, naphthol-AS-BI-phosphohydrolase and α-glucosidase.

The phenotypic characteristics of strain CCPDSM^T^ were compared with those available from *G. greenwoodii*. Distinguishing features are summarised in Table 3.

**Table 3.**
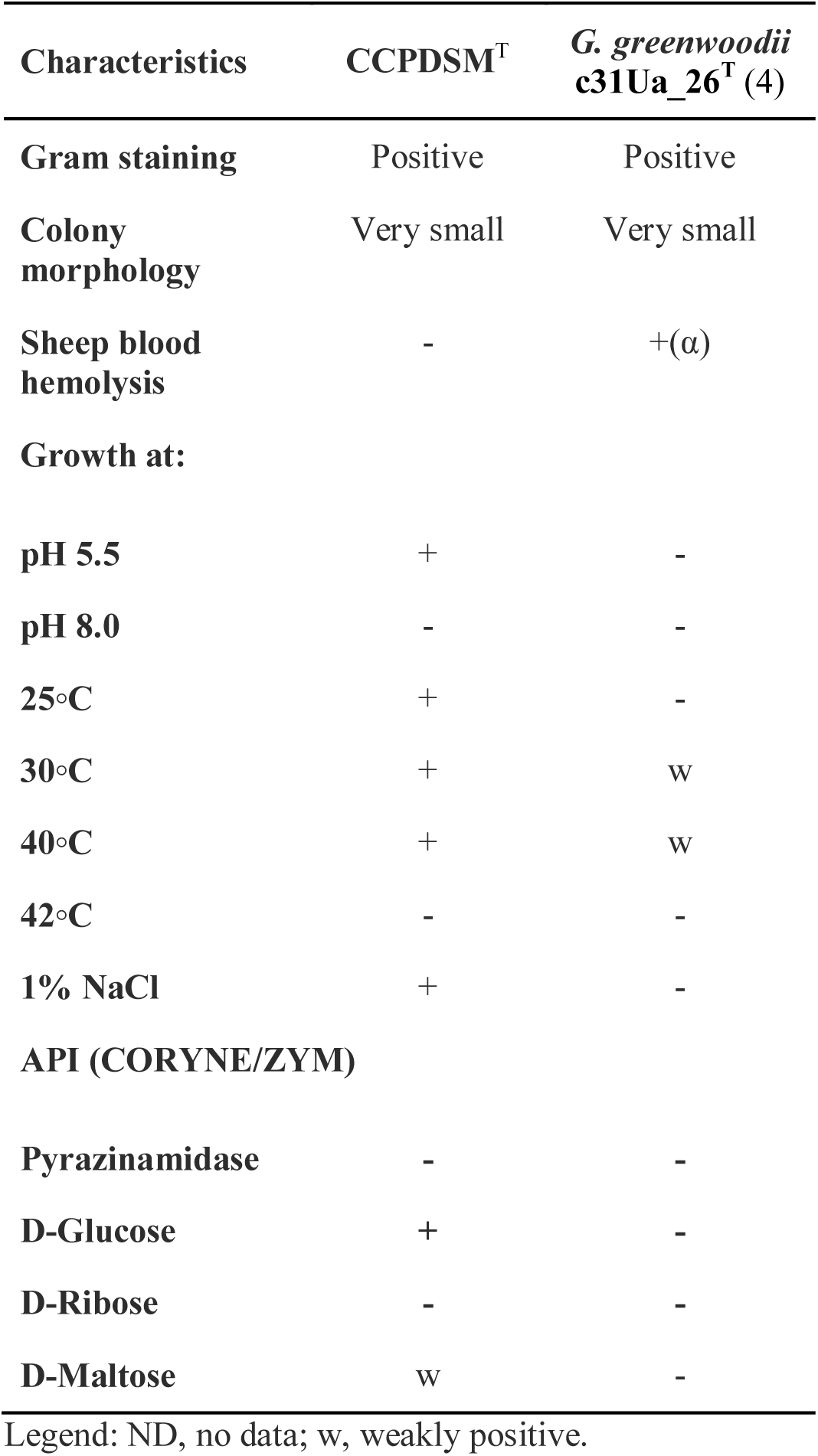
Differential phenotypic characteristics of strain CCPDSM^T^ and its closest relative *G. greenwoodii*.

| Characteristics | CCPDSM<sup>T</sup> | *G. greenwoodii*<br>c31Ua_26<sup>T</sup> (4) |
| --- | --- | --- |
| **Gram staining** | Positive | Positive |
| **Colony morphology** | Very small | Very small |
| **Sheep blood hemolysis** | - | +(α) |
| **Growth at:** |  |  |
| **pH 5.5** | + | - |
| **pH 8.0** | - | - |
| **25°C** | + | - |
| **30°C** | + | w |
| **40°C** | + | w |
| **42°C** | - | - |
| **1% NaCl** | + | - |
| **API (CORYNE/ZYM)** |  |  |
| **Pyrazinamidase** | - | - |
| **D-Glucose** | + | - |
| **D-Ribose** | - | - |
| **D-Maltose** | w | - |

Legend: ND, no data; w, weakly positive.

**Table 4.**
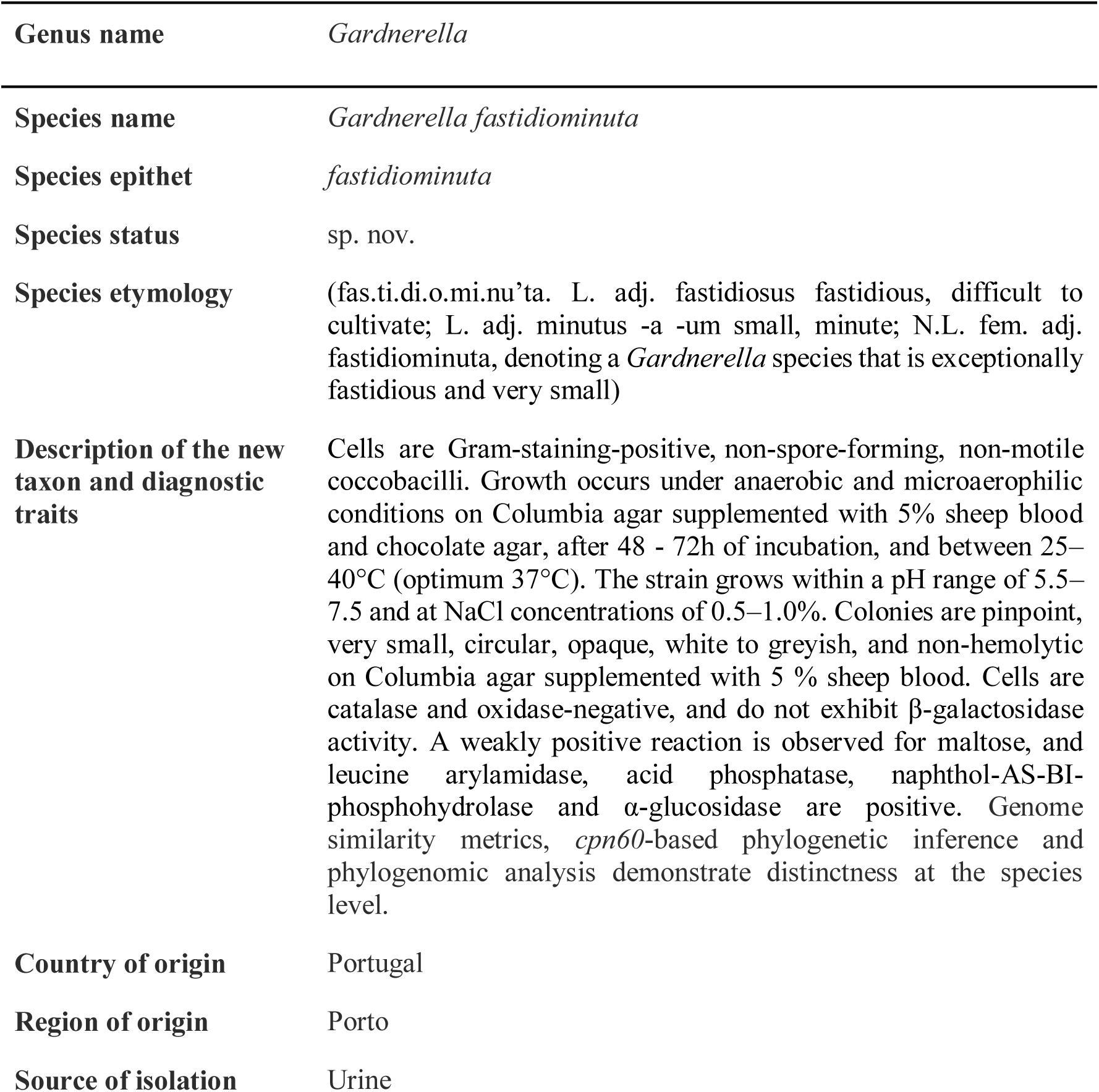
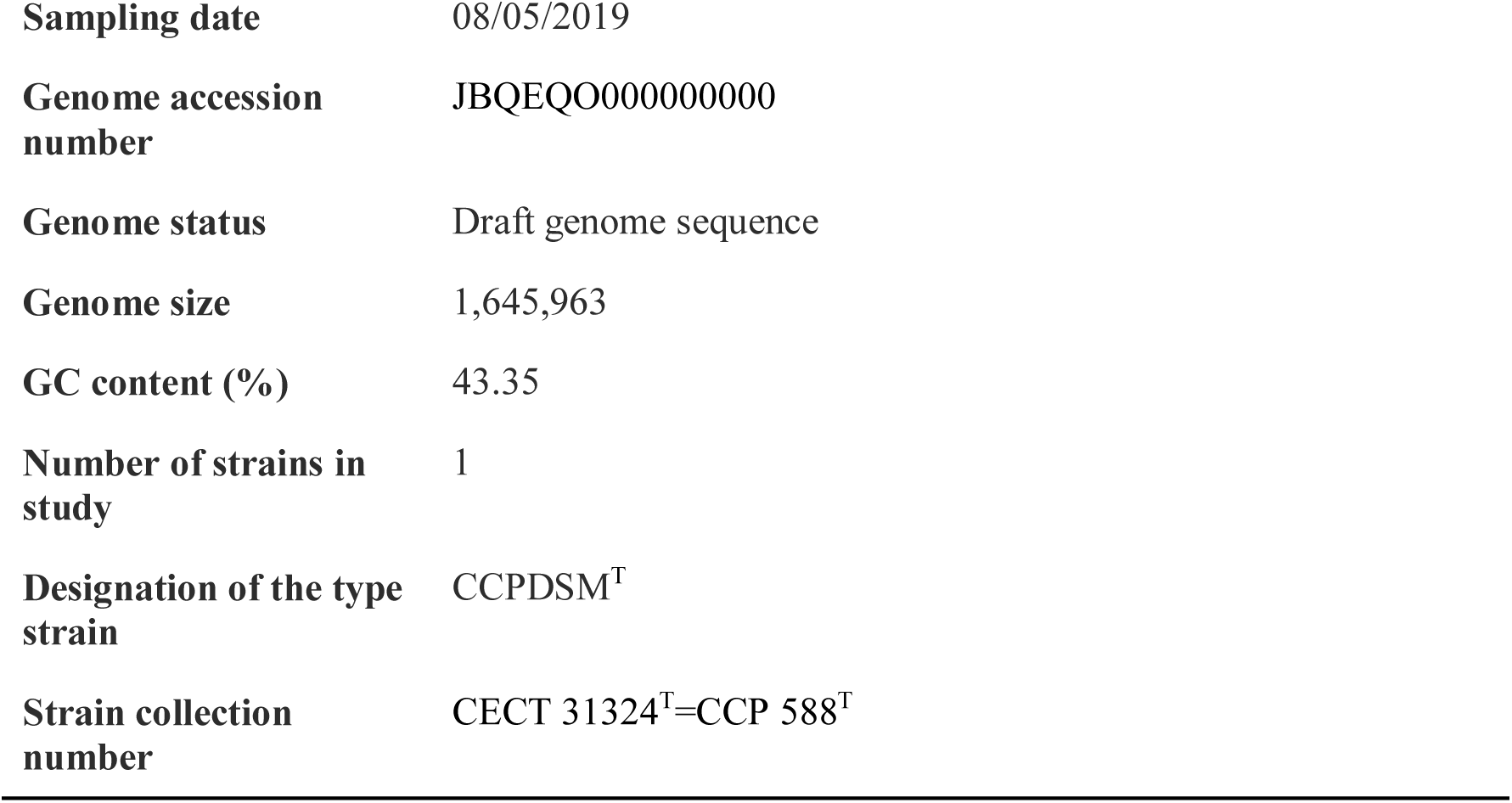
Description of *Gardnerella fastidiominuta* sp. nov.

### Predicted functional comparative genomics

Genome annotation revealed a bacterial strain with a compact metabolic repertoire consistent with a heterotrophic lifestyle and limited biosynthetic autonomy. The genome comprises 1,275 protein-coding CDSs with functional annotation, placing it toward the upper range among the type strains of validly published *Gardnerella* species (from 1,124 in *G. greenwoodii* to 1,328 in *G. massiliensis*) (Table 1). A total of 350 unique enzymes were predicted, alongside 81 transporters and 13 secretion-related genes. The abundance of predicted transporters (6.4% of all annotated protein-coding CDSs) suggests a strong dependence on the uptake of exogenous substrates, in line with adaptation to a nutrient-variable host-associated environment.

Analysis of central carbon metabolism indicated the presence of only a partial set of enzymes associated with carbohydrate processing and downstream metabolic pathways. Notably, enzymes involved in pyruvate conversion and acetyl-CoA metabolism were identified, including pyruvate kinase (EC 2.7.1.40), phosphoenolpyruvate carboxylase (EC 4.1.1.31), phosphate acetyltransferase (EC 2.3.1.8) and acetate kinase (EC 2.7.2.1), as well as acetaldehyde dehydrogenase (EC 1.2.1.10). These enzymes support the potential conversion of pyruvate into acetyl-CoA and subsequently into acetate, indicating the capacity for substrate-level phosphorylation and the production of acetate as a possible metabolic end product.

However, several key enzymes required for a complete Embden–Meyerhof–Parnas pathway were not identified, including phosphofructokinase, fructose-bisphosphate aldolase, glyceraldehyde-3-phosphate dehydrogenase, and phosphoglycerate kinase. The absence of these enzymes does not support the presence of a complete canonical glycolytic pathway. Instead, the metabolic network appears to rely on partial pathways and possibly on the uptake and processing of intermediate metabolites available in the environment. Genes predicted to be involved in starch utilization were identified, suggesting the potential capacity to metabolize complex carbohydrates.

Downstream of pyruvate, the predicted metabolic repertoire is consistent with a fermentative end-product profile. In particular, genes encoding phosphotransacetylase (EC 2.3.1.8) and acetate kinase (EC 2.7.2.1) were identified, supporting potential conversion of acetyl-CoA to acetate via the Pta–AckA pathway, with concomitant ATP generation by substrate-level phosphorylation. Additionally, genes predicted to encode enzymes involved in the conversion of propanoyl-CoA to propionate were identified, suggesting the potential formation of propionate as a fermentation end product. However, the carbon source for propionyl-CoA could not be unambiguously inferred from the annotation data and warrants further investigation. Taken together, these findings support a metabolism compatible with fermentative energy conservation, consistent with the anaerobic and microaerophilic growth phenotype observed and with fermentative traits reported for other members of the genus *Gardnerella* (42).

The genome encodes a complete F₀F₁-ATP synthase complex (EC 3.6.3.14). In the context of the predicted fermentative metabolic profile and the absence of genes encoding cytochrome c oxidases, this complex may function in the ATP hydrolysis direction, consuming ATP to generate a transmembrane proton motive force, as described in fermentative bacteria (43). However, the strain is also capable of growth under microaerophilic conditions, and the genomic basis for this phenotype remains unclear, as no genes encoding canonical terminal oxidases were identified. The presence of redox-related proteins, such as a glutaredoxin-like protein (NrdH), may indicate a limited capacity for intracellular redox homeostasis, which may contribute to tolerance to low oxygen levels.

The genome encodes several enzymes involved in folate metabolism, including dihydrofolate reductase (EC 1.5.1.3), folylpolyglutamate synthase (EC 6.3.2.17), and thymidylate synthase (EC 2.1.1.45), supporting the interconversion and utilisation of folate derivatives required for nucleotide biosynthesis and cellular maintenance. Genes encoding key enzymes of nucleotide salvage pathways were identified, including hypoxanthine-guanine phosphoribosyltransferase (EC 2.4.2.8), adenine phosphoribosyltransferase (EC 2.4.2.7), xanthine phosphoribosyltransferase (EC 2.4.2.22), uracil phosphoribosyltransferase (EC 2.4.2.9), nucleoside hydrolases (EC 3.2.2.1), and uridine phosphorylase (EC 2.4.2.3). These enzymes suggest the potential for active purine and pyrimidine salvage. However, no gene encoding ribose-phosphate pyrophosphokinase (PRPP synthase; EC 2.7.6.1) was identified, indicating that the genomic basis for PRPP supply remains unresolved.

The genome also encodes enzymes of the non-oxidative phase of the pentose phosphate pathway (M00007), including transketolase and transaldolase, supporting the interconversion of sugar phosphates. In contrast, genes corresponding to the oxidative phase of the pathway were not identified, suggesting that NADPH generation via this route is likely absent.

Consistent with the observed phenotypic characterisation, no genes encoding catalase, cytochrome c oxidase, or β-galactosidase were identified in the genome, supporting the catalase-negative, oxidase-negative, and ONPG-negative phenotypes.

Comprehensive genomic analysis revealed the presence of acquired genes associated with antimicrobial resistance. Among these, *lsaC* (91.62% sequence identity, 96% coverage) was identified and has been associated with reduced susceptibility to lincosamides, including clindamycin, a commonly used agent in the treatment of genital tract infections (4). Additionally, the genes *mef(A)* and *msr(D)* (99.84% sequence identity, 98% coverage and 100% sequence identity and 98% coverage, respectively), which are associated with macrolide efflux system, were identified. CRISPR-Cas systems, complete prophages, plasmids, and bacteriocins were not detected.

Overall, genome-based functional prediction depicts a bacterial strain with a reduced but sufficient metabolic network, characterized by carbohydrate-based metabolism, production of short-chain fatty acids, limited biosynthetic autonomy, and extensive transport capabilities. These genomic features are consistent with those reported for members of the genus *Gardnerella* and support its classification within this taxonomic framework.

## Conclusions

In this study, a polyphasic taxonomic approach combining phylogenetic, phylogenomic, genome-based relatedness, phenotypic and functional genomic analyses demonstrated that strain CCPDSMᵀ, isolated from the female urinary microbiome, represents a novel species within the genus *Gardnerella*. Phylogenetic inference based on *cpn60* and phylogenomic analysis consistently placed CCPDSMᵀ as a distinct and well-supported lineage, clearly separated from all validly published *Gardnerella* species.

Genome-wide relatedness metrics confirmed species-level divergence from all *Gardnerella* type strains, while supporting the assignment of closely related publicly available genomes to the same taxon. Phenotypic characteristics and genome-based functional predictions further distinguished CCPDSMᵀ from its closest relatives and were congruent with traits previously reported for members of the genus, while also revealing features consistent with a fastidious, fermentative lifestyle adapted to the urogenital niche.

Together, these data provide robust evidence for the recognition of *Gardnerella fastidiominuta* sp. nov., expanding the known diversity of the genus *Gardnerella* and further highlighting the female urinary tract as a reservoir of previously unrecognized *Gardnerella* species. The formal description of this species contributes to ongoing efforts to refine *Gardnerella* taxonomy and provides a framework for future ecological and clinical investigations into the role of distinct *Gardnerella* species in the urogenital microbiome.

## Credit authorship contribution statement

Liliane P. Ferrador: Methodology, Formal analysis, Investigation, Writing – original draft. Bárbara Duarte: Investigation, Data curation, Strain preservation. Filipa Grosso: Writing – review & editing. Luísa Peixe: Writing – review & editing, Validation, Supervision, Resources, Project administration, Conceptualization. Teresa G. Ribeiro: Methodology, Investigation, Writing – original draft, Writing – review & editing, Validation, Supervision, Conceptualization.

## Acknowledgements

The authors are grateful to Dr. Magdalena Ksiezarek for sample processing.

